# Quantifying the Risk and Cost of Active Monitoring for Infectious Diseases

**DOI:** 10.1101/156497

**Authors:** Nicholas G Reich, Justin Lessler, Jay K Varma, Neil M Vora

## Abstract

During outbreaks of deadly emerging pathogens (e.g., Ebola, MERS-CoV) and bioterror threats (e.g., smallpox), actively monitoring potentially infected individuals aims to limit disease transmission and morbidity. Guidance issued by CDC on active monitoring was a cornerstone of its response to the West Africa Ebola outbreak. There are limited data on how to balance the costs and performance of this important public health activity. We present a framework that estimates the risks and costs of specific durations of active monitoring for pathogens of significant public health concern. We analyze data from New York City’s Ebola active monitoring program over a 16-month period in 2014-2016. For monitored individuals, we identified unique durations of active monitoring that minimize expected costs for those at “low (but not zero) risk” and “some or high risk”: 21 and 31 days, respectively. Extending our analysis to smallpox and MERS-CoV, we found that the optimal length of active monitoring relative to the median incubation period was reduced compared to Ebola due to less variable incubation periods. Active monitoring can save lives but is expensive. Resources can be most effectively allocated by using exposure-risk categories to modify the duration or intensity of active monitoring.

## Introduction

The outbreak of Ebola virus disease (Ebola) in west Africa was the largest outbreak of a highly virulent acute infection in modern times, leading the global health community to reassess the ability of highly pathogenic acute infections to spread widely in today’s interconnected world. To improve rapid identification and evaluation of individuals infected with Ebola, on October 27, 2014, the U.S. Centers for Disease Control and Prevention (CDC) recommended active monitoring of individuals potentially exposed to Ebola virus. Individuals under active monitoring were asked to contact local health authorities to report their health status every day for 21 days after their last potential exposure.

Thousands of individuals were monitored for Ebola in the United States between October 2014 and February 2016, including over 10,000 individuals during one five month period.(1) CDC’s guidance on active monitoring was a cornerstone of its response to Ebola. The guidance balanced numerous stakeholder concerns, including mitigating risk to communities and travelers without unnecessarily restricting individual liberties. Recommendations for active monitoring were discontinued in February 2016.(2) Over 20% of all individuals actively monitored for Ebola in the United States were monitored in New York City (NYC), more than any other jurisdiction.(1)

Active monitoring may help prevent and contain outbreaks of rapidly spreading emerging pathogens that pose a grave threat to public health. Such outbreaks may occur naturally, via a bioterrorist attack, or via unintended release from a laboratory. The decision to implement active monitoring depends on an assessment of the risk posed by a pathogen and the ability of active monitoring to reduce that risk. Key considerations include the transmissibility and pathogenicity of the pathogen, the potential size of an outbreak, and the relationship between the time of symptom onset and infectiousness.(3) Ebola, Middle East Respiratory Syndrome Coronavirus (MERS-CoV), and smallpox are examples of viral illnesses for which active monitoring could play a pivotal role in preventing a large-scale outbreak.(4)

Active monitoring programs must balance conflicting priorities. The central goals of these programs are to identify, isolate, and treat infected individuals quickly. Setting an active monitoring period many times longer than any known incubation period of the pathogen of interest could virtually guarantee that all infected individuals would exhibit symptoms while being monitored. However, such a program would be unreasonably expensive, inconvenience monitored individuals, and incur many financial and social costs through frequent responses to false positive cases. Evidence-based monitoring periods and appropriate tailoring of the monitoring intensity to disease risk should therefore be used to balance costs with biosecurity risks.

Here we present an empirical framework for evaluating the risks and costs associated with active monitoring (implemented in an online tool available at http://iddynamics.jhsph.edu/apps/shiny/activemonitr/). We apply this framework to Ebola, MERS-CoV, and smallpox using data on the natural history of these diseases and data from the Ebola response of the NYC Department of Health and Mental Hygiene (DOHMH).

## Methods

### Estimating the incubation period distribution

The incubation period of a disease is the duration of time between exposure to the pathogen and symptom onset.(5) This characteristic is imperfectly observed in most settings.(6,7) We obtained previously published incubation period observations on 145 cases of Ebola in Guinea (8), 170 laboratory-confirmed cases of MERS-CoV in South Korea(9) and 362 cases of smallpox(10–12). We fit parametric distributions to the observed incubation period data using maximum likelihood techniques (see Supplemental Text). We ran sensitivity analyses to evaluate the influence of several outlying observations in the Ebola dataset.

### A model for active monitoring outcomes

We developed a model that uses varying probabilities to alter the incubation period, thus estimating whether active monitoring would identify an individual infected with the pathogen of interest. Figure 1 presents the model schema, which involves a monitored individual having one of four outcomes:

1. no symptoms warranting clinical follow-up,
2. no symptomatic infection with the disease of interest, but occurrence of symptoms that necessitate ruling out of the disease of interest,
3. symptomatic infection with the disease of interest occurring during the individual’s period of active monitoring, and
4. symptomatic infection with the disease of interest occurring outside the individual’s period of active monitoring.

**Figure 1.**
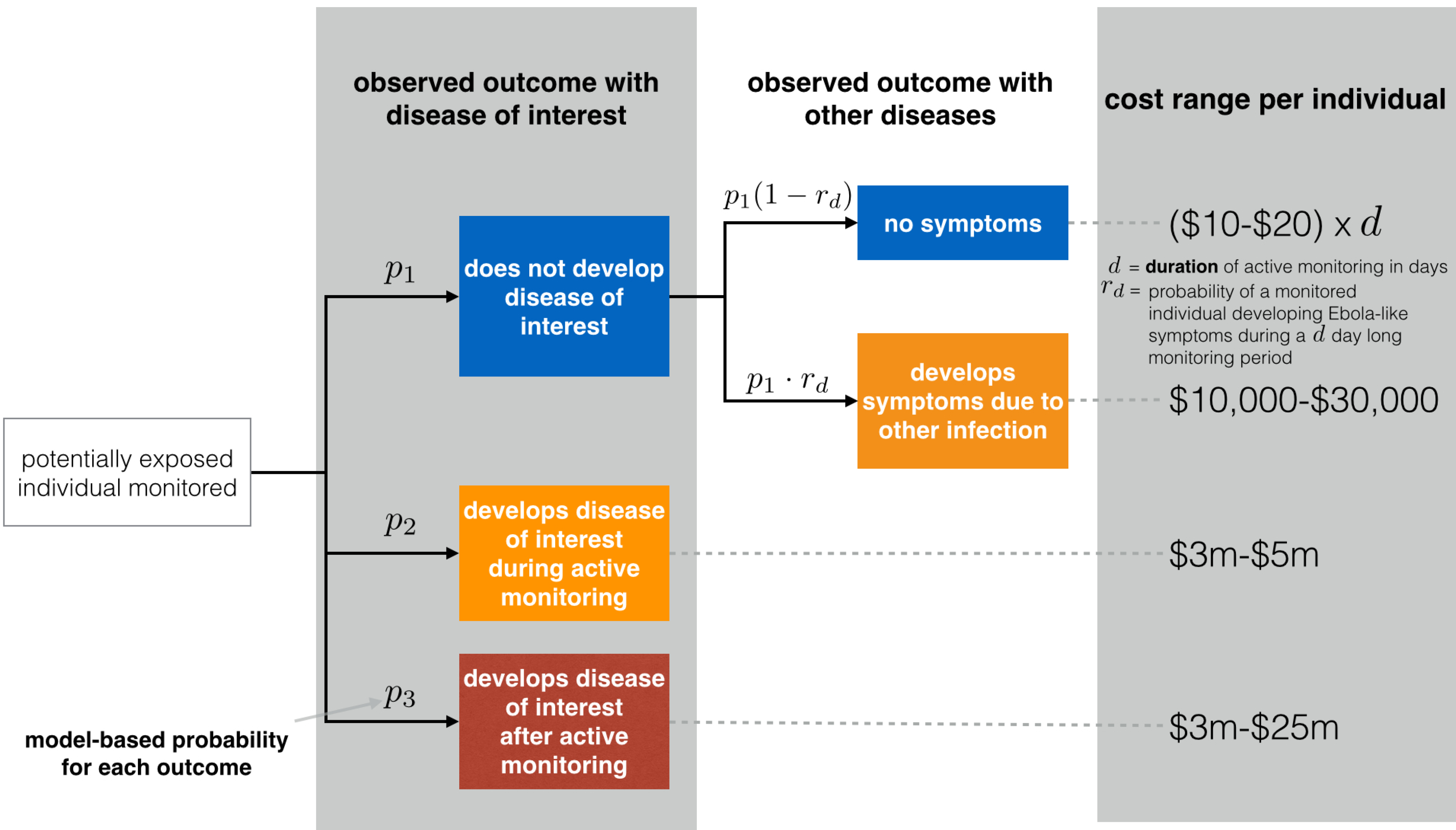
Model schematic representing four outcome scenarios for a person under active monitoring. Estimated costs shown are based on published costs(23) as well as new data from DOHMH and Bellevue Hospital in NYC. Details on the model formulation are available in Supplemental Table 3. Our model uses probabilities to calculate the likelihood of each of the possible model outcomes. Additionally, we estimate the probability that an individual who does not develop the disease of interest develops symptomatic illness necessitating hospitalization to rule-out the disease of interest. (See Supplemental Text for details.)

Scenario 4 represents the highest risk of secondary transmission and may also reduce the chance of a positive clinical outcome for the sick individual.

Using this model, we calculate the probability of each of the four outcomes, with the associated expected costs, by combining the data on the probabilities of infection, the estimated incubation period distribution, and plausible cost ranges for different outcomes. Since limited data prevent us from formally estimating variance, we created conservative (i.e. maximally wide) intervals for possible costs by using the endpoints of the plausible cost ranges. Additionally, it determines the uncertainty associated with these estimates due to not having precise incubation period observations and having an unknown time between exposure and the beginning of monitoring for a particular individual (see Supplemental Text). For these pathogens, we assume that infectiousness coincides with the onset of symptoms.(13–15)

Our model estimates the risks and costs associated with active monitoring programs for a range of active monitoring durations. To estimate the cost per person-day of monitoring, we used data on the number of individuals actively monitored by DOHMH and costs associated with the DOHMH Ebola response. Additionally, for the purposes of hypothetical cost calculations, we assumed that an individual who becomes symptomatic with the disease of interest while under active monitoring gives rise to no secondary infections, while an individual who develops symptoms after his/her active monitoring period ends could give rise to as many as 4 new Ebola infections (an upper estimate based on prior research (16,17)).

We developed open-source software, including a freely-available web application at http://iddynamics.jhsph.edu/apps/shiny/activemonitr/. The source code for the web app, the data for the analyses, and the code to reproduce this manuscript itself are all freely available online under an open-source license at GitHub (https://github.com/reichlab/activemonitr), with a static version in an open-access digital library (18). All analyses were run in **R** version 3.3.1 (2016-06-21).(19) These tools enable others to easily implement our model and reproduce our results.

### Stratifying by exposure risk

Classifying individuals based on prior exposure risks enables targeted strategies in a range of public health response settings, including active monitoring. For example, in response to the West Africa Ebola outbreak, CDC issued recommendations on risk stratification of individuals for a potential Ebola virus exposure (“high risk”, “some risk”, “low (but not zero) risk”, “no identifiable risk”) and for how long and how intensively individuals in each of these categories should be monitored.(20) DOHMH’s active monitoring program, described previously(21), was implemented consistent with CDC recommendations. However, creating, evaluating, and modifying such classifications in practice is a difficult task and requires situational awareness and data that would vary depending on the pathogen and outbreak setting.

For the CDC Ebola risk strata, we estimated probabilities of a monitored individual developing Ebola. These estimates were based on extrapolated numbers of actively monitored individuals in the United States during 2014-2016(1) and public data on the four domestic cases of Ebola(22) (Supplemental Text, Table 2).

## Results

### Incubation period estimates

Estimates and credible regions for incubation period distribution parameters for Ebola, MERS-CoV, and smallpox are shown in Figure 2. Consistent with other studies that used the same data,(8,9,12) we estimated that half of all cases of Ebola will have an incubation period of less than 8.9 days (95% CI: 8.0-9.8), of MERS-CoV less than 6.9 days (95% CI: 6.3-7.5) and of smallpox less than 12.2 days (95% CI: 12.0-12.4). Additionally, the data suggest that 95% of cases of Ebola will have an incubation period of less than 20.3 days (95% CI: 18.1-23.0), of MERS-CoV 13.3 days (95% CI: 12.0-14.8), and of smallpox 15.8 days (95% CI: 15.4-16.2).

**Figure 2.**
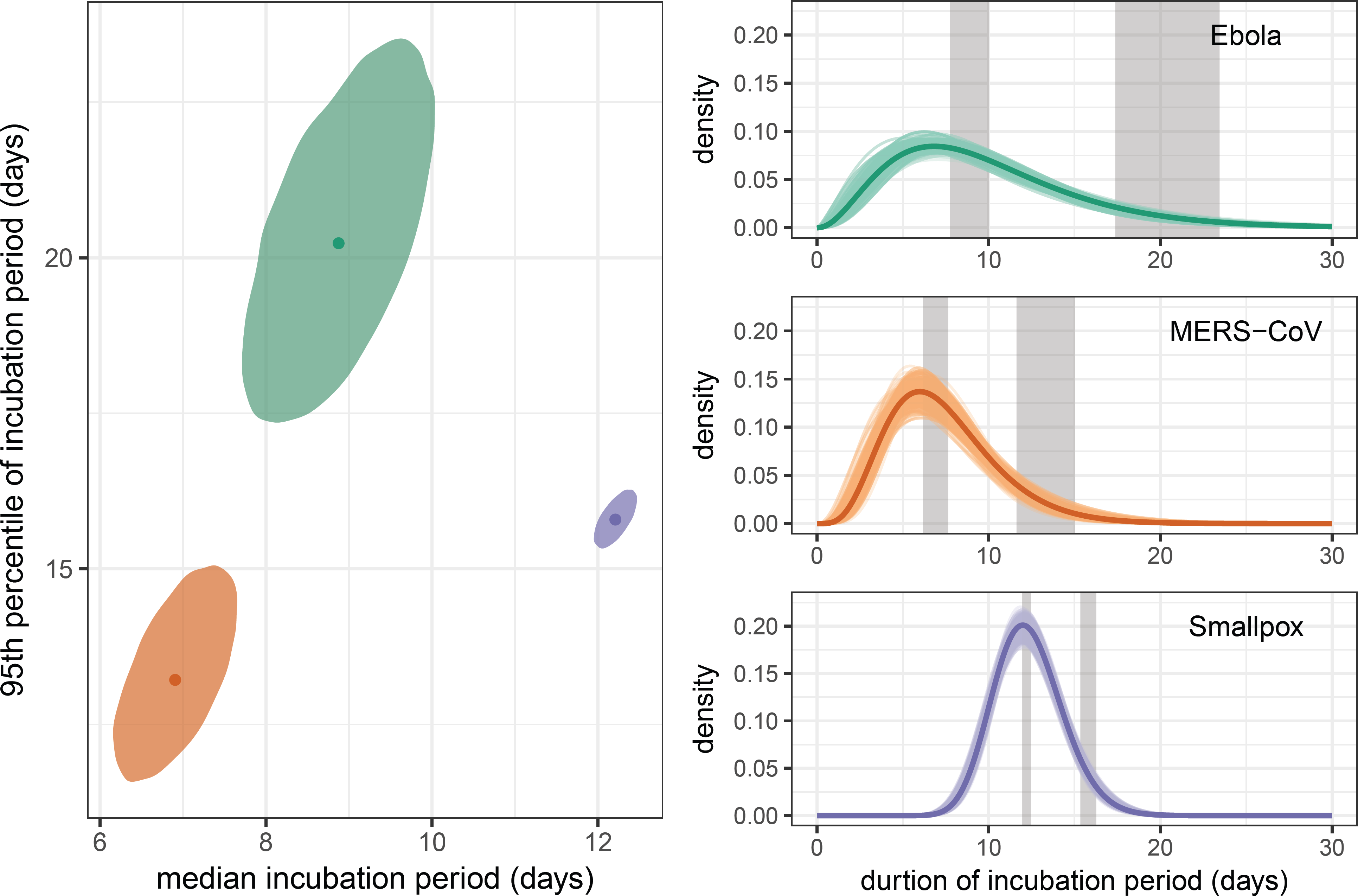
Estimates and credible regions for incubation period distributions for Ebola, MERS-CoV and smallpox. The shaded elliptical areas represent regions that contain 95% of the estimated posterior distributions for each of the three diseases. The disease-specific curves plotted on the right show the estimated distribution for the incubation period for each disease (dark line). To show some of the uncertainty associated with these estimates, a random selection of density functions sampled from the joint posterior are represented by colored transparent lines around the heavy lines. Shaded vertical bands indicate the marginal credible regions for the median and 95th percentile.

We estimated smallpox to have the longest median incubation period of the three pathogens considered, although its distribution also showed the least overall variance. We estimate that MERS-CoV incubation periods are the shortest of the three, with the upper limit of the 95th percentile being just above 15 days (Figure 2).

### Modeling the risk of symptomatic illness

Using information on cases of Ebola diagnosed in the U.S., as well as extrapolated case information, we estimated the probability of a “some-risk” or “high-risk” individual developing a symptomatic infection with Ebola as 1 in 1,000. For “low (but not zero) risk” individuals, we estimated this risk to be 1 in 10,000 (Supplemental Text, Table 2).

Incorporating these probabilities into our model, for each pathogen we calculated the probability of symptomatic illness developing in an individual after they were no longer being actively monitored. To standardize the results across diseases, we present the risk estimates across a range of different active monitoring period durations, shown as multiples of the median incubation period for each disease (Figure 3).

**Figure 3.**
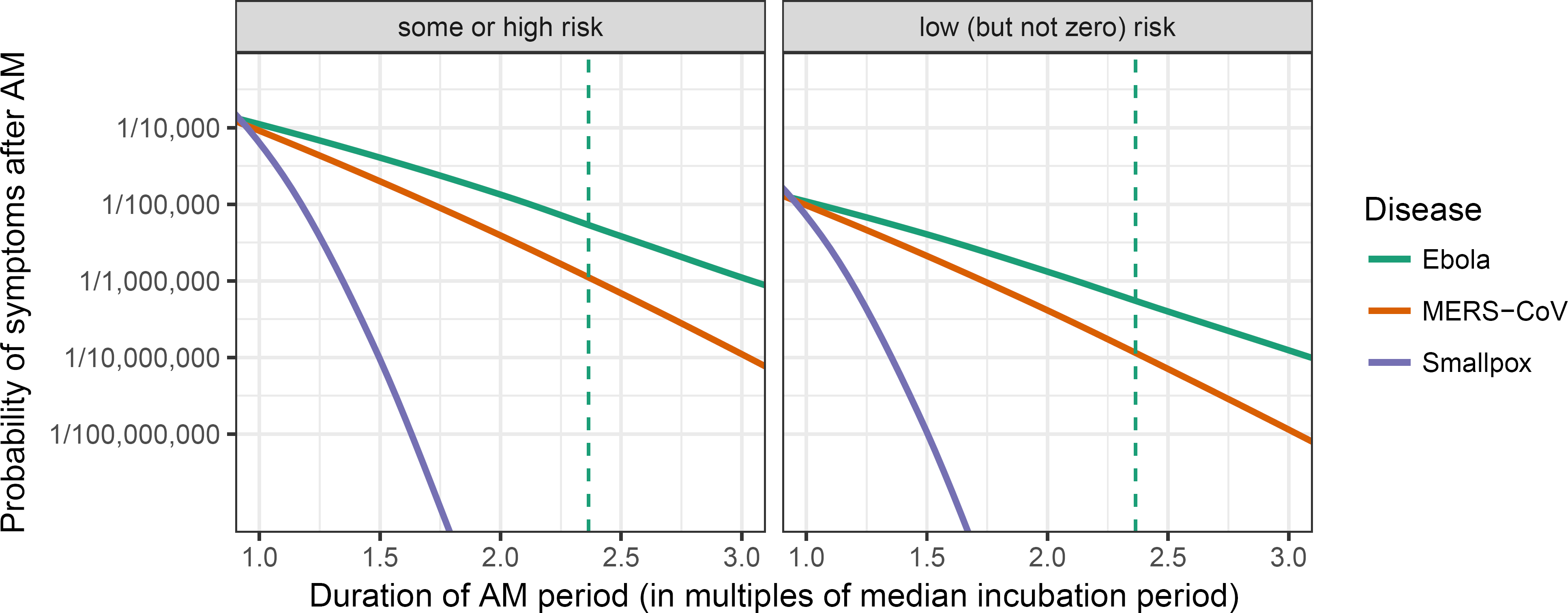
Estimated probabilities of symptoms occurring after active monitoring (AM) ends across different active monitoring period durations, shown as multiples of the median incubation period. Figures are shown for ‘some or high risk’ and ‘low (but not zero) risk’ scenarios, with probabilities of developing symptomatic infection set to 1/1,000 and 1/10,000, respectively. The vertical dashed line indicates the 21 day duration recommended for Ebola active monitoring (i.e., approximately 2.4 times the median incubation period of Ebola).

The model showed that an increase in the duration of the active monitoring period is associated with a decline in the probability that an individual develops symptoms due to the disease of interest after the active monitoring period ends. The rate at which that risk declines depends on the estimated variability of the incubation period distribution. Since the estimated incubation period distribution for Ebola showed the highest variability, the probability of symptoms occurring after active monitoring ends decreased more slowly when compared with the other pathogens. This feature also impacts the sensitivity of estimating the optimal duration of active monitoring when the probabilities of developing symptomatic infection are mis-specified (Supplemental Text, Table 4).

During the West Africa Ebola outbreak, CDC recommended active monitoring for 21 days. This corresponds to about 2.4 times the median incubation period for Ebola (vertical dashed line, Figure 3). To achieve the same level of absolute risk for either MERS-CoV or smallpox, our model suggests that a duration of active monitoring would need to be set at less than two times the median incubation period for both of these diseases.

### Minimizing costs of active monitoring programs: Ebola case study

The total expense for Ebola response by DOHMH during the period from July 31, 2014-November 7, 2015 was $9.7 million. Of this, $4.3 million was in response to a single Ebola case (23) and $1.9 million was for active monitoring. The remaining balance of $3.5 million was used for other Ebola preparedness activities. DOHMH monitored 5,379 non-unique individuals during this period (active monitoring began in NYC on October 25, 2014). We used these data to estimate that the cost of monitoring per person-day in NYC was $10–$20.

The number of serious infection events occurring in actively monitored individuals in NYC was small. None of the monitored individuals developed Ebola. The single Ebola case diagnosed in NYC occurred before active monitoring was implemented. We assumed that a small fraction of monitored individuals, 30 (or 0.6%), developed symptoms within the 21 day monitoring period that were serious enough to necessitate hospitalization at an institution in NYC to rule-out Ebola. At one hospital, Bellevue, it cost between $10,000-$30,000 per hospitalization to rule out Ebola in symptomatic, actively monitored individuals (John Maher, NYC Health and Hospitals, personal communication). This does not include the substantial infrastructural costs of creating and maintaining a special pathogens unit at the hospital.

Our model provides ranges of expected cost for active monitoring systems. It identifies an optimal duration of active monitoring, by finding the expected cost range with the lowest maximum value. We applied the model to the case-study of Ebola in NYC, based on data from DOHMH (Table 1, Figure 4). Expected costs of short periods of active monitoring (left hand side of Figure 4) are driven by the cost of a missed case and the number of expected additional secondary cases, while the rate of decline with additional days of monitoring is driven by the shape of the incubation period distribution. The costs of longer periods of active monitoring are driven by the per day cost of monitoring and costs of false positive detections (right hand side of Figure 4).

**Figure 4.**
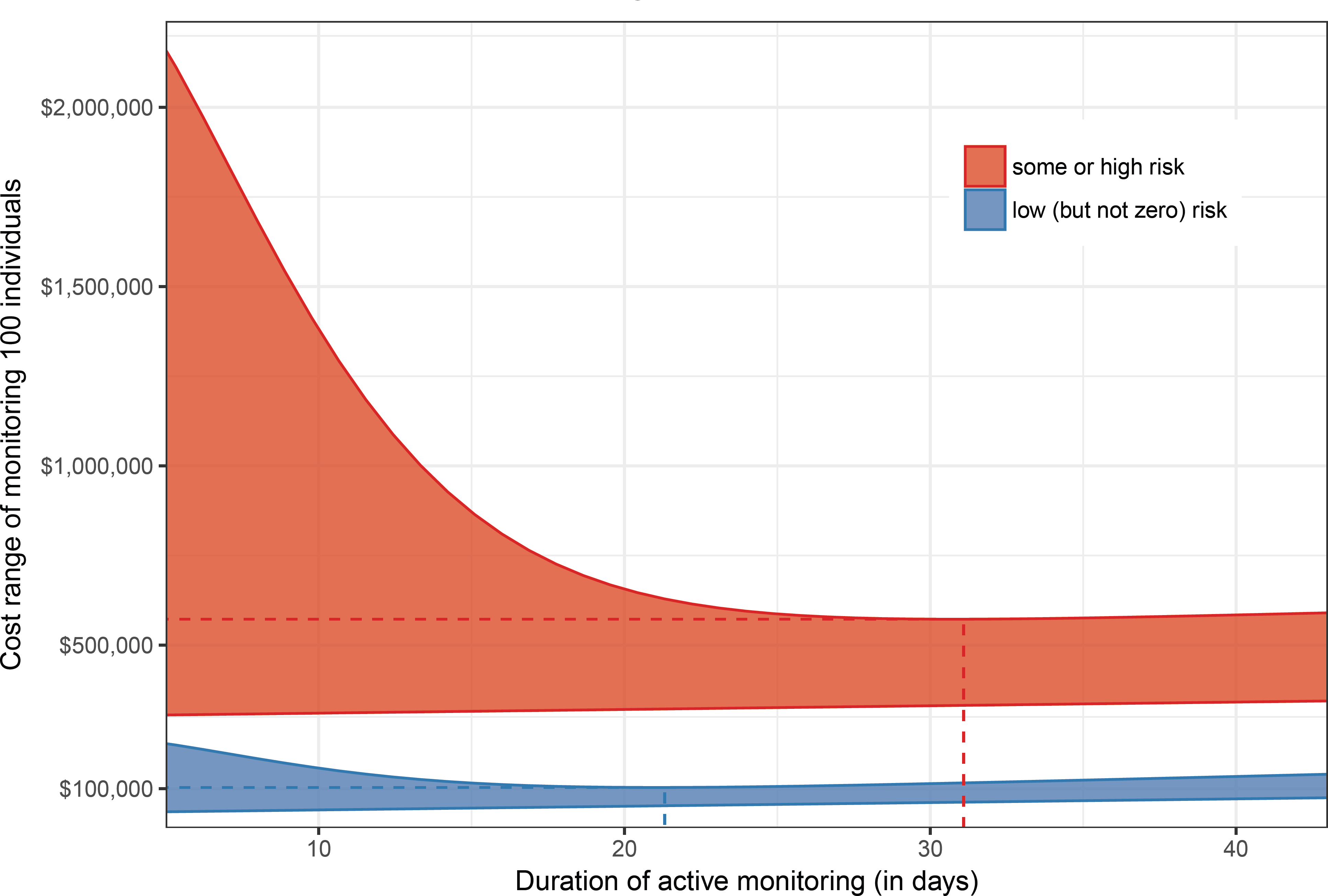
Estimated cost ranges of actively monitoring 100 individuals for Ebola, calculated separately for some or high risk individuals and low (but not zero) risk individuals. The dashed lines intersect at the minimum point for the upper limit of each cost range.

**Table 1:** Total expected costs to the public health system (including active monitoring and response) per 100 monitored individuals, at two exposure-risk categories. Columns represent multiples of the median incubation period of Ebola. Rows represent the CDC exposure-risk category. Costs are inclusive of active monitoring and public health response and are based on model outputs for Ebola. Values are in $’000s.

|  | 1.5 | 2 | 3 | 5 |
| --- | --- | --- | --- | --- |
| some risk and high risk | \$314-\$1,002 | \$318-\$726 | \$327-\$579 | \$346-\$593 |
| low (but not zero) risk | \$44-\$125 | \$48-\$106 | \$57-\$108 | \$76-\$143 |

For the low-risk individuals, the model suggests that the cost is minimized with 21.3 days of monitoring (i.e., 2.4 median incubation periods, 95% CI = 19.5 - 22.2). For the some- or high-risk individuals, the model suggests that the cost is minimized with 31.1 days of monitoring (i.e., 3.5 median incubation periods, 95% CI = 27.5 - 33.8) (Figure 4). The model results are sensitive to the assumed cost per day of monitoring and the number of secondary cases. For example, if the upper limit of the cost increases from $20 to $40/day or the number of secondary cases decreases from 4 to 2, the durations of monitoring that would minimize expected cost decrease by 2-3 days for each scenario.

## Discussion

For three pathogens of significant public health concern, we quantified the risk, costs, and attendant uncertainty associated with specific durations of active monitoring. The approach presented can be used to develop empirically based public health policies in future outbreaks.

Perception of risk is a key component of decision making, but ultimately science should inform public health policies. A key challenge faced by decision makers is that the public’s perception of risk is often determined by the consequences of the event even if the probability of the event occurring is rare. With severe diseases, the consequences may be perceived as so dire that the public’s tolerance for risk will be very low. While no policy can guarantee zero risk, analyses such as those presented here can assist in determining appropriate thresholds that reduce risk below a level tolerable for a risk-averse population.

In some settings, active monitoring could serve as an alternative to quarantine, which involves both monitoring and severe restrictions on movement. In countries at low risk for Ebola, quarantine raised many ethical issues during the West Africa Ebola outbreak.(24) CDC’s Ebola active monitoring recommendations were therefore a critical cornerstone of the public health response in the United States that strived to balance the concerns from a diverse set of stakeholders without unnecessarily restricting individual liberties.(2) Active monitoring has since been applied by DOHMH for an individual at risk for another acute viral illness, Lassa fever. However, for diseases in which asymptomatic persons may transmit infection –such as influenza or measles– detection of the cases based on active monitoring of symptomology might not be soon enough to prevent secondary transmission. As such, for these diseases, quarantine might be a more appropriate public health measure.

Our model suggests durations of active monitoring for emerging pathogens early in an outbreak. Accurate estimates of the median incubation period can be made even when few data are available.(6) While accurately estimating the variability of the incubation period requires more data(25), existing data on the variation observed in similar pathogens could be used to inform early estimates until better data become available. When estimating incubation periods in real-time, care must be taken to adjust for possible truncation of longer incubation periods (26,27) or selection biases that could favor reporting of shorter incubation periods (9).

While our model explicitly propagates uncertainty in the incubation period distribution, public health practitioners will usually have too little data to estimate uncertainty for other model parameters. To help account for this limitation, we ran the model for plausible ranges of other parameters, presenting a range of possible costs and risks. Sensitivity to these and other assumptions can be assessed through the open-access web applet that implements the model.

Evaluating the cost of monitoring programs is a challenging, multi-faceted problem. While we present new data from New York City’s active monitoring program showing that the per-individual cost of monitoring is “low” (about $10-$20 per day), the total program cost can be substantial. Limiting who needs to be monitored and the duration of monitoring could provide valuable savings.

Some cost-efficiency may be achieved through targeted strategies. As CDC recommended for Ebola, it may be appropriate to use exposure-risk categories to modify the duration or intensity of active monitoring. Additionally, stronger public health messaging could encourage travelers to take necessary health precautions to avoid common travel-associated illnesses, e.g. malaria.(28) This, in turn, could reduce unnecessary testing and costs associated with symptomatic events with diseases other than the one of interest. Low-cost versions of active monitoring have the potential to add value to public health response to less severe outbreaks as well.

Accurately estimating the risk of developing disease within exposure-risk categories poses a challenge to public health officials. Our analyses show that the sensitivity to mis-specified risk varies by pathogen. When the estimated cost varies across plausible risk levels, the full range of potential costs and durations should be considered until better data are available to obtain more precise estimates of the risk.

The framework presented is generally applicable, but given the lack of empirical cost information for the response to and monitoring of smallpox and MERS-CoV our analysis is strongly focused on Ebola. While we have used some of the best available data on large-scale active monitoring programs in our analyses, some potential costs and savings were not factored in to these calculations. The estimates available from DOHMH do not include the costs of cell phones given to monitored individuals, the staff time incurred by CDC and US Department of Homeland Security in screening travelers at airports, and the costs of evaluating individuals with symptoms that were not due to Ebola. More detailed cost data was not available from DOHMH, but such data could help to disentangle the fixed costs associated with running an active monitoring program from the cost per monitored individual. Additionally, we only presented gross cost estimates for the evaluation of a symptomatic individual under active monitoring from one NYC hospital, which was not representative of all hospitals in NYC. Finally, local considerations prompted DOHMH to opt for an active monitoring program involving humans to make and receive active monitoring reports from individuals. In contrast, other jurisdictions used digital systems to administer monitoring reports.(29) The merits of these different approaches deserve further investigation.

Recent history has shown that the unexpected emergence of new disease threats has become a recurring theme in global health and preparedness. While active monitoring does not play a role in the response to every emerging infectious disease (e.g., it has not played an important role in the response to the Zika virus epidemic), it will likely be used again in the response to future threats. Our framework provides valuable information for assessing the cost-effectiveness of various active monitoring strategies in response to critical disease outbreaks. By providing an empirical basis for evaluating active monitoring programs, these tools can strengthen biosecurity and optimize active monitoring programs in response to future global disease threats.

## Acknowledgments

We thank the individuals who underwent active monitoring in NYC, Ifeoma Ezeoke for preparing the DOHMH active monitoring data and Margaret Pletnikoff for preparing the DOHMH cost data. We thank Dr. Simon Cauchemez, Dr. Ousmane Faye, and Dr. Amadou Sall for providing access to the Ebola incubation period data from Guinea and giving permission for this data to be made publicly available. We also acknowledge John Maher for providing details on costs of evaluating Ebola patients at Bellevue Hospital in New York City. NGR was funded by a grant from the MIDAS program of the National Institutes of General Medical Sciences (R35GM119582). The findings and conclusions in this article are those of the authors and do not necessarily represent the official position of DOHMH, CDC, NIGMS, or the National Institutes of Health.

## Bibliography

1. Stehling-Ariza T, Fisher E, Vagi S, Fechter-Leggett E, Prudent N, Dott M, et al. Monitoring of Persons with Risk for Exposure to Ebola Virus Disease - United States, November 3, 2014-March 8, 2015.MMWR Morbidity and mortality weekly report. 2015 Jul;64(25):685–9.

2. Centers for Disease Control and Prevention. Interim U.S. guidance for monitoring and movement of persons with potential Ebola virus exposure [Internet]. Available from: http://www.cdc.gov/vhf/ebola/exposure/monitoring-and-movement-of-persons-with-exposure.html

3. Fraser C,Riley S, Anderson RM, Ferguson NM. Factors that make an infectious disease outbreak controllable. Proceedings of the National Academy of Sciences of the United States of America. 2004 Apr;101(16):6146–51.

4. Centers for Disease Control and Prevention. Middle East Respiratory Syndrome: People at Increased Risk for MERS [Internet]. 2015. Available from: http://www.cdc.gov/coronavirus/mers/risk.html

5. Brookmeyer R. Incubation Period of Infectious Diseases. In: Encyclopedia of biostatistics. Chichester, UK: John Wiley & Sons, Ltd; 2005.

6. Reich NG, Lessler J, Cummings DAT, Brookmeyer R. Estimating incubation period distributions with coarse data. Statistics in medicine. 2009 Sep;28(22):2769–84.

7. Lessler J, Reich NG, Brookmeyer R, Perl TM, Nelson KE, Cummings DAT. Incubation periods of acute respiratory viral infections: a systematic review. The Lancet Infectious Diseases. 2009 May;9(5):291–300.

8. Faye O, Boëlle P-Y, Heleze E, Faye O, Loucoubar C, Magassouba N, et al. Chains of transmission and control of Ebola virus disease in Conakry, Guinea, in 2014: an observational study. The Lancet Infectious Diseases. 2015 Mar;15(3):320–6.

9. Virlogeux V, Park M, Wu JT, Cowling BJ. Association between Severity of MERS-CoV Infection and Incubation Period. Emerging Infectious Diseases. 2016;22(3).

10. Litvinjenko S, Arsic B, Borjanovic S. Epidemiologic Aspects of Smallpox in Yugoslavia in 1972. Bulletin of the World Health Organization. 1972 Nov;

11. Mack TM. Smallpox in Europe, 1950-1971. Journal of Infectious Diseases. 1972 Feb;125(2):161–9.

12. Nishiura H. Determination of the appropriate quarantine period following smallpox exposure: an objective approach using the incubation period distribution. International journal of hygiene and environmental health. 2009 Jan;212(1):97–104.

13. Velásquez GE, Aibana O, Ling EJ, Diakite I, Mooring EQ, Murray MB. Time from Infection to Disease and Infectiousness for Ebola Virus Disease, a Systematic Review. Clinical infectious diseases: an official publication of the Infectious Diseases Society of America. 2015 Oct;61(7):1135–40.

14. Cowling BJ, Park M, Fang VJ, Wu P, Leung GM, Wu JT. Preliminary epidemiological assessment of MERS-CoV outbreak in South Korea, May to June 2015. Euro Surveillance. 2015;20(25):7–13.

15. Fenner F, Henderson DA, Arita I, Jezek Z, Ladnyi ID. Smallpox and its Eradication. World Health Organization; 1988.

16. Althaus CL. Estimating the Reproduction Number of Ebola Virus (EBOV) During the 2014 Outbreak in West Africa. PLoS currents. 2014;6.

17. Chowell G, Hengartner NW, Castillo-Chavez C, Fenimore PW, Hyman JM. The basic reproductive number of Ebola and the effects of public health measures: the cases of Congo and Uganda. Journal of Theoretical Biology. 2004 Jul;229(1):119–26.

18. Reich NG, Li X. reichlab/activemonitr: preprint. Zenodo [Internet]. 2017; Available from: https://zenodo.org/record/260135

19. R Core Team. R: A Language and Environment for Statistical Computing. Vienna, Austria: R Foundation for Statistical Computing; 2015.

20. Centers for Disease Control and Prevention. Epidemiologic Risk Factors to Consider when Evaluating a Person for Exposure to Ebola Virus [Internet]. 2015. Available from:http://www.cdc.gov/vhf/ebola/exposure/risk-factors-when-evaluating-person-for-exposure.html

21. Millman AJ, Chamany S, Guthartz S, Thihalolipavan S, Porter M, Schroeder A, et al. Active Monitoring of Travelers Arriving from Ebola-Affected Countries - New York City, October 2014-April 2015. MMWR Morbidity and mortality weekly report. 2016;65(3):51–4.

22. Centers for Disease Control and Prevention. Cases of Ebola Diagnosed in the United States | Ebola Hemorrhagic Fever | CDC [Internet]. Available from: http://www.cdc.gov/vhf/ebola/outbreaks/2014-west-africa/united-states-imported-case.html

23. Yacisin K, Balter S, Fine A, Weiss D, Ackelsberg J, Prezant D, et al. Ebola virus disease in a humanitarian aid worker - New York City, October 2014. MMWR Morbidity and mortality weekly report. 2015 Apr;64(12):321–3.

24. Drazen JM, Kanapathipillai R, Campion EW, Rubin EJ, Hammer SM, Morrissey S, et al. Ebola and quarantine. The New England journal of medicine. 2014 Nov;371(21):2029–30.

25. World Health Organization. Consensus document on the epidemiology of severe acute respiratory syndrome (SARS) [Internet]. 2003. Available from: http://aje.oxfordjournals.org.silk.library.umass.edu/content/163/3/211.full.pdf+html

26. Brookmeyer R, Blades N, Hugh-Jones M, Henderson DA. The statistical analysis of truncated data: application to the Sverdlovsk anthrax outbreak. Biostatistics. 2001 Jun;2(2):233–47.

27. Nishiura H, Inaba H. Estimation of the incubation period of influenza A (H1N1–2009) among imported cases: addressing censoring using outbreak data at the origin of importation. Journal of Theoretical Biology.2011 Mar;272(1):123–30.

28. Ezeoke I, Saffa A, Guthartz S, Tate A, Varma JK, Vora NM. Health Precautions Taken by Travelers to Countries with Ebola Virus Disease. Emerging Infectious Diseases. 2016 May;22(5):929–31.

29. Parham M, Edison L, Soetebier K, Feldpausch A, Kunkes A, Smith W, et al. Ebola active monitoring system for travelers returning from West Africa—Georgia, 2014–2015. MMWR Morbidity and mortality weekly report. 2015 Apr;64(13):347–50.

